# Pathogenic *ASXL1* somatic variants in reference databases complicate germline variant interpretation for Bohring-Opitz Syndrome

**DOI:** 10.1101/090720

**Authors:** Colleen M. Carlston, Anne H. O’Donnell-Luria, Hunter R. Underhill, Beryl B. Cummings, Ben Weisburd, Eric V. Minikel, Daniel P. Birnbaum, Exome Aggregation Consortium, Tatiana Tvrdik, Daniel G. MacArthur, Rong Mao

**Author notes:** Correspondence should be addressed to: Colleen M. Carlston, PhD, Department of Pathology, University of Utah, 15 North Medical Drive East, Salt Lake City, UT, USA 84112. C.M. Carlston and A.H. O’Donnell-Luria contributed equally to this work.

## Abstract

The interpretation of genetic variants identified during clinical sequencing has come to rely heavily on reference population databases such as the Exome Aggregation Consortium (ExAC). Genuinely pathogenic variants, particularly in genes associated with severe autosomal dominant conditions, are assumed to be absent or extremely rare in these databases. Clinical exome sequencing of a six-year-old female patient with seizures, global developmental delay, dysmorphic features and failure to thrive identified an *ASXL1* variant that was previously reported as causative of Bohring-Opitz syndrome (BOS). Surprisingly, the variant was observed seven times in the ExAC database, presumably in individuals without BOS. Although the BOS phenotype matched the presentation of the patient, the presence of the variant in reference population databases introduced ambiguity in result interpretation. Interrogation of the literature revealed that acquired somatic mosaicism of *ASXL1* variants (including known pathogenic variants) during hematopoietic clonal expansion may be concomitant with aging in healthy individuals. We examined all high quality *ASXL1* predicted truncating variant calls in the ExAC database and determined the majority could be attributed to this phenomenon. Failure to consider somatic mosaicism may lead to the inaccurate assumption that conditions like Bohring-Opitz syndrome have reduced penetrance, or the misclassification of potentially pathogenic variants.

## INTRODUCTION

The pace of discovery of genes underlying inherited conditions has accelerated, primarily due to decreasing costs and improved access to next-generation sequencing technology (Boycott et al. 2013). Unfortunately, in some cases misattribution of disease-association to genetic variants has occurred when these variants were under-sampled in a control set of unaffected individuals (Andreasen et al. 2013). The advent of large, openly accessible population databases has helped address this issue (MacArthur et al. 2014). Guidelines for variant assessment incorporate population frequency data; absence from population controls is considered moderate evidence for pathogenicity (Richards et al. 2015). If the population frequency of a variant vastly exceeds the incidence of the associated condition, the variant would typically be considered benign, or at least only weakly penetrant (Richards et al. 2015).

Bohring-Opitz syndrome (BOS) is a rare, autosomal dominant, malformation syndrome, characterized by severe intrauterine growth restriction, poor feeding, hypotonia, profound intellectual disability associated with brain MRI findings, prominent metopic suture, trigonocephaly, micro/retrognathia, exophthalmos, nevus flammeus of the face that fades with age, low-set ears, low hairline, upslanting palpebral fissures, broad alveolar ridges, and hirsutism (Hoischen et al. 2011; Magini et al. 2012; Dangiolo et al. 2015; Russell et al. 2015; Urreizti et al. 2016). A frequent finding is the “BOS posture”, characterized by flexion of the elbows and wrists with deviation of the wrists and metacarpophalangeal joints. Previously BOS patients often succumbed to recurrent infections, but better clinical management has resulted in more patients surviving to adulthood (Hoischen et al. 2011; Russell et al. 2015). Two patients have been reported with early onset of pubertal changes (Russell et al. 2015). Screening for Wilms tumors is also recommended for children diagnosed with BOS (Russell et al. 2015).

BOS is caused by *de novo* truncating variants in the *additional sex combs-like 1* (*ASXL1*) gene (Hoischen et al. 2011), which encodes a putative member of the Polycomb Repressive Complex (Fisher et al. 2003). The molecular mechanism of *ASXL1* pathogenic variants in BOS is unknown, but the C-terminal PHD (zinc finger domain), implicated in mediating interactions with other proteins, may be lost when ASXL1 protein is truncated due to premature stop codons. This has been proposed to ablate the auto-inhibition of ASXL1, which is consistent with observed over-activation of the mutant ASXL1-BAF1 complex (Balasubramani et al. 2015). In contrast, the DNA-binding domain at the N-terminus of ASXL1 is predicted to remain intact. Sometimes the same pathogenic variants are found in myeloid malignancies and in BOS (Hoischen et al. 2011; Genovese et al. 2014; Jaiswal et al. 2014), and similar mechanisms of pathogenesis may occur in both conditions. Several BOS-associated pathogenic variants occur in the last exon of *ASXL1* (Hoischen et al. 2011; Magini et al. 2012; Boycott et al. 2013) and would not be predicted to trigger nonsense mediated decay. *Asxl1* heterozygous null mice exhibit craniofacial dysmorphism with reduced penetrance (Abdel-Wahab et al. 2013), though reduced penetrance has never been observed with Bohring-Opitz syndrome in humans. The following analysis will refer to putative protein truncating variants as pathogenic when they are known to cause BOS or truncating if there is no current data on pathogenicity.

## MATERIALS AND METHODS

### Clinical case report

The proband presented to our clinic at 6 years of age with a history of seizures and developmental delay. She was the fifth child (four healthy older siblings) born at term via normal spontaneous vaginal delivery after an uncomplicated pregnancy to a 30-year-old mother and a 34-year-old father. Parents were non-consanguineous. Her initial difficulty with feeding, hypotonia, and an unusual cry precipitated genetic testing for Cri-du-chat syndrome that was negative. Her subsequent course included seizures (tonic seizures presented at <1 month of age; tonic-clonic seizures developed by 2 years of age), nearly absent corpus callosum, dysphagia (G-tube dependent since 1 year of age), profound developmental delay, and a distinct facial appearance.

Physical exam identified a small child with length, weight, and head circumference below the first percentile. She had bilateral ptosis, optics for her myopia, a faint nevus flammeus of the face, protuberant cheeks, high palate with a narrow jaw and bruxism, tented lips and an open mouth, diffuse hirsutism including mild synophrys, and mild fat pads on her metacarpophalangeal joints. She was non-verbal and her eyes did not track light but rather roamed continuously. She was able to reach and grab objects and bring them to her mouth, but was unable to sit independently. The characteristic BOS posture was not present.

The patient had an extensive genetic evaluation at multiple institutions. The following testing was performed: karyotype (46, XX), SNP-microarray, methylation studies for Prader-Willi/Angelman syndrome, *SMN1* dosage for spinal muscular atrophy, and sequencing for *FOXG1*, *MECP2*, *UBE3A*, *PNPO*, and *ALDH7A1* genes, lysosomal enzyme activity, very long-chain fatty acids, CSF neurotransmitters, transferrin isoelectric focusing for congenital disorders of glycosylation, and screening for mucopolysaccharidosis. All studies were normal.

### Exome capture enrichment

Genomic DNA from peripheral blood was extracted using a Gentra PureGene Blood Kit (Qiagen, Germantown, MD) and DNA concentration was quantified with a NanoDrop 8000 (Thermo Scientific, Wilmington, DE). DNA (3 µg) was sheared to 180 base pair (bp) fragments. Illumina adapters were added using SureSelect XT kit reagents (Agilent Technologies, Santa Clara, CA). Adapter-ligated DNA were hybridized to exome-targeted biotinylated RNA baits (SureSelect Clinical Research Exome, Agilent Technologies, Santa Clara, CA) for 24 hours at 65 °C. Hybridized DNA were captured with streptavidin-coated magnetic beads. Exome targets of interest were eluted and then barcoded/indexed. DNA quality was assessed using a LabChip Gx (Perkin Elmer, Waltham, MA), and samples were pooled.

### Sequencing

Samples were sequenced on a HiSeq2500 platform (Illumina, San Diego, CA) using paired-end 100 bp reads. Sequences were aligned to the human genome reference (GRch37) using Burrows-Wheeler Aligner (BWA 0.5.9) (Li and Durbin 2009) with default parameters. PCR duplicates were removed with Samtools (Li et al. 2009), and base quality score recalibration, local realignment, and variant calling were performed with Genome Analysis Toolkit (GATK v1.3) (McKenna et al. 2010). Parental variants were filtered to identify *de novo* variants, which were then confirmed by Sanger sequencing.

### Exome Aggregation Consortium (ExAC) analysis

We reviewed high-quality putative truncating variants (nonsense, frameshift, and canonical splice-disrupting variants) in *ASXL1* present in ExAC (Lek et al. 2016). High-quality sites met the following criteria: (1) they were given a PASS filter status by VQSR (GATK v3.1 Haplotype Caller algorithm), (2) the depth (DP) of the read coverage was >=10 and the genotype quality (GQ) was >=20. IGV screenshots (Thorvaldsdóttir et al. 2013) were reviewed along with variant position in Ensembl transcripts on the UCSC Genome Browser (Kent et al. 2002). This data, including IGV readviz of supporting read data, is available at http://exac.broadinstitute.org/ or for download at http://exac.broadinstitute.org/downloads (including the non-TCGA ExAC subset). For TCGA samples, sequence data from germline samples only (not tumor samples) is included in ExAC. Age data, which were available for a subset of ExAC, have been rounded to the closest 5-year mark with a minimum of 20 and maximum of 85 for privacy. To assess variant balance for the *ASXL1* gene, we also evaluated rare non-coding or in-frame (non-frameshift) indels (allele count <10, consistent with frequency of truncating variants), along with 30 randomly selected (using a random number generator) rare (allele counts <10) synonymous or missense variants in *ASXL1*. Only heterozygous variants were included in this analysis and to determine allele balance the *ASXL1* gene region was recalled from BAM files. The allele balance data from ExAC was not retained so large-scale evaluation of allele balance is not possible in ExAC. Allelic imbalance was defined as fewer than 35% of sequencing reads derived from the variant allele, as this represents roughly two standard deviations below the mean for synonymous variants (95% confidence interval). We also evaluated whether variants were listed in ClinVar (Supplementary Table 1) (Landrum et al. 2016).

## RESULTS

Exome sequencing performed on the patient, both parents, and two unaffected siblings, identified a *de novo* variant, c.1210C>T; p.Arg404Ter, in the *ASXL1* gene (NM_015338). When examined in IGV, the *ASXL1* variant had high-quality reads inconsistent with a sequencing artifact, which was further supported by Sanger sequencing (Supplementary Figure 1).This variant has been previously reported to cause BOS (Hoischen et al. 2011) and is listed in the dbSNP database (rs373145711) as pathogenic referencing the ClinVar entry (variant ID 30986) (Landrum et al. 2016). The p.Arg404Ter variant is also listed as a disease-causing mutation for BOS in HGMD^®^(Stenson et al. 2003). However, this variant was also reported in the Exome Aggregation Consortium (ExAC) browser (7/121378 chromosomes, with no homozygotes) and in the Exome Variant Server (2/13006 chromosomes, with no homozygotes) (Tennessen et al. 2012) (although one of these samples is present in both datasets).

The presentation of the patient closely resembles the description of BOS and the p.Arg404Ter variant had been previously reported in another BOS patient (Hoischen et al. 2011). Furthermore, all reported BOS patients (including this patient) that have *de novo ASXL1* truncating variants are severely affected, and decreased penetrance or variable expressivity has not been described in BOS. Therefore, the presence of this variant in these reference population databases was incongruous with the expectation of pathogenicity for this variant.

We explored the possibility that the observation of the *ASXL1* p.Arg404Ter variant in a population database might be the result of somatic mosaicism. Hematopoietic clonal expansion of cells with *ASXL1* pathogenic variants has been reported as a frequent event acquired with aging (Genovese et al. 2014; Jaiswal et al. 2014; Xie et al. 2014). So-called clonal hematopoiesis of indeterminate potential (CHIP) is present in roughly 10% of individuals over age 65 and nearly 20% of those over age 90 (Genovese et al. 2014; Jaiswal et al. 2014; Steensma et al. 2015). The most commonly mutated genes in CHIP are *DNMT3A*, *TET2*, and *ASXL1*; all three encode epigenetic modifiers. About 1% of patients with somatic variants in these genes progress each year to be diagnosed with myelodysplastic syndrome (MDS), a group of clonal hematologic stem cell disorders characterized by ineffective hematopoiesis and an increased risk of developing acute myeloid leukemia. Although somatic *ASXL1* variants likely contribute to malignancy (Genovese et al. 2014; Jaiswal et al. 2014), the detection of pathogenic *ASXL1* variants alone is not considered diagnostic of MDS given the low rate of progression.

In contrast, germline, *de novo ASXL1* variants have been convincingly shown to cause BOS (Hoischen et al. 2011). Therefore, it is important to distinguish between germline and somatic *ASXL1* variants. The gold standard to prove somatic mosaicism is to test DNA derived from other tissue sources such as buccal swabs or hair follicles, but due to the need to protect the anonymity of those who contribute their genetic information to reference databases further testing of these individuals is generally not possible. However, manual examination of the read support for seven individuals in ExAC who share our patient’s p.Arg404Ter *ASXL1* variant revealed that five of the seven demonstrate considerable allelic imbalance (fewer than 35% of sequencing reads derived from the variant allele) (Table 1). Of the two individuals with a variant allele balance close to 50%, one was an individual with cancer (variant present in 43% of reads) and the other was an 85-year-old woman (variant present in 42% of reads). As discussed above, *ASXL1* truncating variants are associated with an increased cancer risk and aging. In summary, two of the seven individuals carrying the p.Arg404Ter variant reportedly have cancer, and the others have a median age of 70 (the youngest is 55 and the oldest 85). Though DNA from other tissues in these individuals is not available for testing, it is possible to infer that the incidence of the p.Arg404Ter *ASXL1* variant in the ExAC database is likely due to somatic mosaicism rather than reduced penetrance of BOS with germline pathogenic variants.

**Table 1:** Bohring-Opitz syndrome-associated *ASXL1* variants

| Variant | Allele Balance | Phenotype | Sex | Age |
| --- | --- | --- | --- | --- |
| p.Arg404Ter | 23% | - | Male | 55 |
| p.Arg404Ter | 42% | - | Female | 85 |
| p.Arg404Ter | 19% | - | Male | 80 |
| p.Arg404Ter | 21% | - | Male | 60 |
| p.Arg404Ter | 43% | Cancer | Female | NA |
| p.Arg404Ter | 13% | Cancer | Male | NA |
| p.Arg404Ter | 12% | - | Male | 70 |
| p.Arg965Ter | 21% | - | Female | 85 |

NA = not available

The observation of the *ASXL1* p.Arg404Ter variant in the ExAC database raised the possibility that other pathogenic *ASXL1* variants might also be found in reference population databases. For example, another BOS-associated pathogenic variant, p.Arg965Ter (Magini et al. 2012), was present in ExAC with an allele balance of 21% in an 85-year-old woman (Table 1). In collaboration with ExAC investigators, all putative truncating *ASXL1* variants in the ExAC database were examined (Supplementary Table 1). A total of 342 individuals were carriers of 56 truncating *ASXL1* variants, all of which were present in a heterozygous state. Four variants (observed in five individuals) were excluded because they were unlikely to cause *ASXL1* truncation for various reasons, including being associated with non-constitutive exons (p.Phe6CysfsTer62 and c.141-2A>G), and a complex mutational event involving two indels in close proximity in the same two individuals (p.Ile1329SerfsTer121 and p.Pro1330ThrfsTer41) whose overall effect was a Ile to Ser substitution rather than a frameshift (Supplementary Figure 2).

The two most common truncating variants, p.Gly646TrpfsTer12 and p.Gly645ValfsTer58, identified in 132 and 118 individuals respectively, were also excluded (Supplementary Figure 2). These variants are located in an eight-nucleotide homopolymer tract of guanines (one variant is 7G and the other 9G) and potentially represent PCR artifacts. Intriguingly, deep sequencing of a large series of myeloid malignancies found the p.Gly646TrpfsTer12 variant (confirmed in several cases by capillary electrophoresis or Sanger sequencing) to be the most common cancer-associated *ASXL1* variant (identified in 47 out of 133 myeloid malignancies) (Van Ness et al. 2016). So far p.Gly645ValfsTer58 has not been observed even once in roughly 2000 myeloid malignancy panels (W. Shen, personal communication). Since polymerase slippage would be expected to create an equal number of 7G and 9G variants, Van Ness et al. conclude that the absence of p.Gly645ValfsTer58 variants suggests p.Gly646TrpfsTer12 by comparison represents a genuine mutational hotspot. Our own analysis potentially supports this assessment, as p.Gly646TrpfsTer12 (median allele balance 21.8%) was associated with TCGA samples in 66% of cases compared to 8% for p.Glys645ValfsTer58 (median allele balance 15.4%). However, despite acknowledging that a subset of the p.Gly646TrpfsTer12 variants are likely genuine, without deeper coverage or additional confirmation (e.g. Sanger sequencing) it would be difficult to distinguish the real variants from the artifacts. For this reason we excluded all 250 individuals with either variant from our analysis.

The remaining 88 individuals (with 50 different truncating *ASXL1* variants) were investigated to determine the relative balance of variant alleles. An IGV view of the mapped reads is publically available on the ExAC browser for up to 5 randomly chosen heterozygous (and up to 5 homozygous when present) individuals for each variant in ExAC. The median variant allele balance was 25% (range 9-59%, interquartile range (IQR) 17.5-34.5%). Since individuals with cancer might be expected to show a higher balance of the *ASXL1* variants, we separated the 23 individuals from the TCGA cohort and the 65 individuals from the non-cancer cohort (from across multiple ancestries and cohorts in ExAC) and compared the variant allele balance in both groups. Since the aim of ExAC was to include germline samples, individuals with hematological cancers were excluded; however, increased rates of hematopoietic mosaicism involving variants in *ASXL1* is also seen in individuals with other cancer types (Artomov et al. 2016). The variant allele balance was not significantly different between the cancer (median 31%) and the non-cancer (median 22%) cohorts (Mann-Whitney U Test, p=0.18). Compared to a median age of 56 for all individual in ExAC, the non-cancer cohort had a median age of 70 (59/65 individuals with associated age), which is potentially concordant with age-related somatic mosaicism (Figure 1).

**Figure 1.**
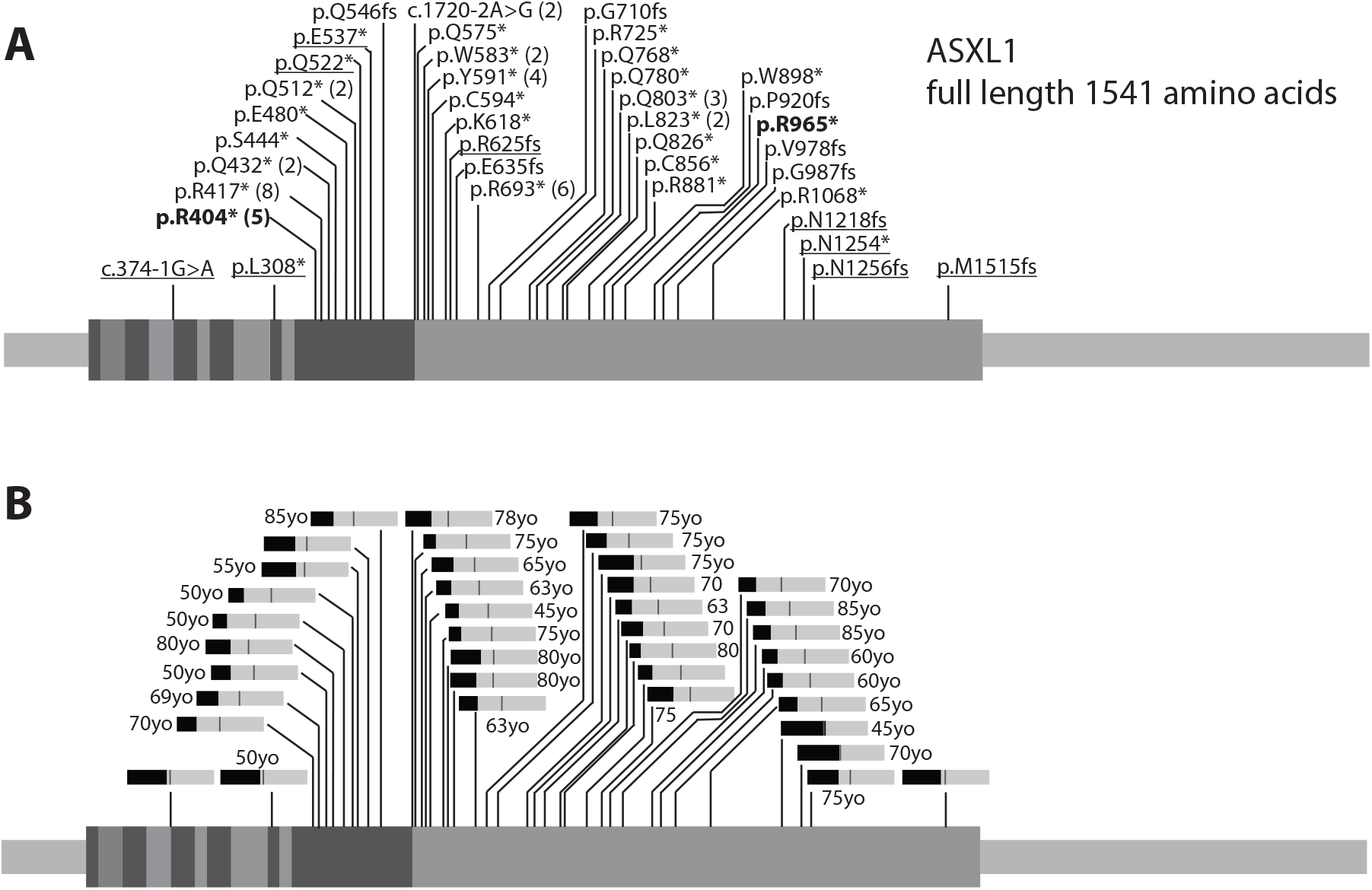
ASXL1 cDNA (exons indicated) and truncating variants after filtering. A) 39 ASXL1 truncating variants (excluding samples from cancer cohorts) observed in Exome Aggregation Consortium browser, with the number of individuals greater than one listed in parentheses (total = 65 individuals). The bolded variants have been observed in patients with Bohring-Opitz syndrome. Underlined variants are those with an average allele balance equal to or greater than 35%. B) Bar graphs of average allele balances for each variant (medium gray bar = 50%), with average ages associated with each variant listed when available. For the original data see Supplementary Table 1.

To ensure that the skewed variant allele balance was not related to sequencing of *ASXL1*, we assessed two other classes of variants that are largely predicted to be neutral in terms of functional impact. First, rare synonymous variants in *ASXL1* had no evidence of variant allele imbalance with a median allele balance of 47% (range 15-61%, IQR 44.3-49.9%). Only three individuals for the 30 synonymous variants analyzed across 70 individuals had an allele balance <35%. The difference between the allele balances for non-cancer-associated, putative truncating variants (median 22%) and for the rare, non-cancer synonymous and missense variants (median 47%) is significant (Mann-Whitney U Test, p=1.02E-14). Second, we were concerned whether difficulty in aligning indels, which constitute 40% of the observed truncating *ASXL1* variants, could contribute to an apparent decreased allele balance. However, there was no evidence of variant allele imbalance in 20 *ASXL1* non-frameshift indels (noncoding and inframe indels) identified in 34 individuals, with a median variant allele balance of 46% (range 29-65%, IQR 40.5-50.0%). Overall, there was no evidence of general allele imbalance for rare *ASXL1* variants that are predicted to not impact protein function.Additionally, we also observed allele imbalance with nonsense variants, which presumably would not be affected by alignment issues.

Sixteen individuals in the non-cancer cohort have putative truncating *ASXL1* variants with allele balances between 35-60%. Seven of these individuals carry variants that were also detected in others with considerable allelic imbalance, suggesting they too may derive from somatic mosaicism. The median age for this group is 75 (age data available for 13 of16 individuals), which is slightly higher than for the non-cancer cohort as a whole, and might hint at an age-related increase in the allele balance of truncating *ASXL1* variants in these individuals. Most of the putative truncating variants are found in the last exon (Figure 1), which is also the largest exon and a location where putative truncating variants are likely to generate transcripts that escape nonsense mediated decay. Comparing the age distribution of the non-cancer cohort with putative truncating *ASXL1* variants (median 70 years old), to the age distribution of the non-cancer cohort with rare synonymous or missense *ASXL1* variants (median 55 years old) reveals a significant shift towards older individuals in the former group (Mann-Whitney U Test, p=3.98E-11) (Figure 2).

**Figure 2.**
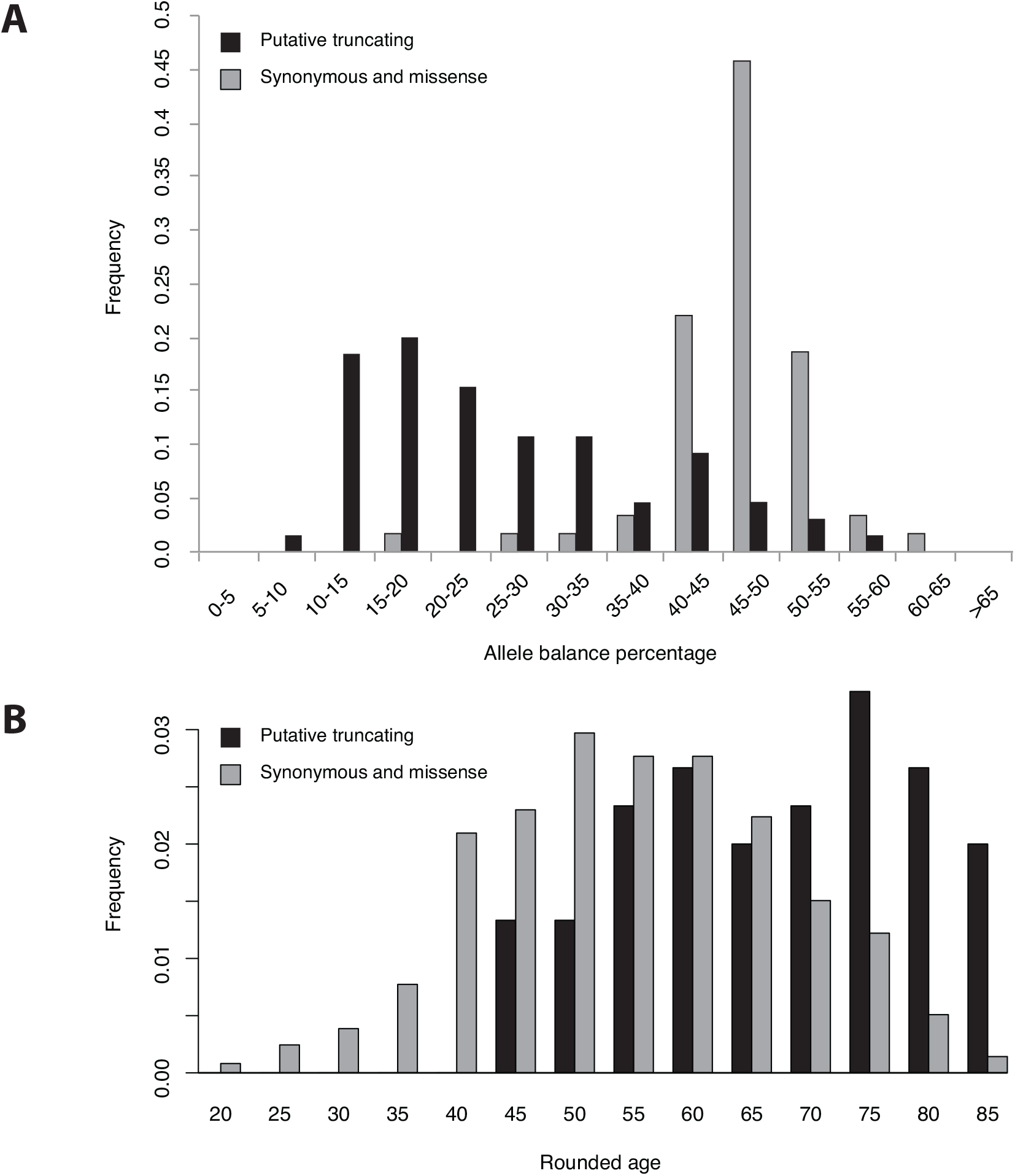
Allele balance and age distribution of individuals from non-cancer cohorts with rare *ASXL1* variants. Black bars indicate individuals with putative truncating *ASXL1* variants and grey bars indicate rare (<10 individuals) synonymous and missense *ASXL1* variants in ExAC. A) The median allele balance of putative truncating *ASXL1* variants was 22% (n=65), while the median allele balance for those with rare synonymous or missense *ASXL1* variants was 47%. The difference in allele balance between these two groups is significant (Mann-Whitney U Test, p=1.02E-14). B) The median rounded age for individuals with putative truncating *ASXL1* variants was 70 (n=60, no age data was available for five individuals), while the median rounded age for those with rare synonymous or missense *ASXL1* variants was 55 (n=983). The age difference between these two groups is significant (Mann-Whitney U Test, p=3.98E-11). For the original data see Supplementary Table 1.

Importantly, there is direct evidence of somatic mosaicism for p.Asn1256MetfsTer24, a variant with an allele balance of 36% present in a single individual in ExAC. This individual also had a common synonymous variant 5 nucleotides upstream of this rare frameshift variant. There were 109 reads covering both variants, 39% of which contained the frameshift deletion alone, 52% the synonymous variant alone, and 9% neither variant (Supplementary Figure 2). Most likely, the frameshift variant arose as a somatic mutation in *trans* to the synonymous variant, although we cannot exclude the less likely possibility that there was partial rescue with loss of the frameshift variant.

In summary, by examining the data associated with 345 individuals cumulatively carrying 56 truncating *ASXL1* variants in the ExAC database, we found evidence suggesting most are unlikely to be germline variants. In a few cases, the variants themselves are suspect (*i.e.* dubious annotation or the homopolymer effect), highlighting the importance of reviewing the variant position on a genome browser. In other cases, allelic imbalance strongly indicated the potential for somatic mosaicism, especially when fewer than 35% of reads were derived from the variant allele, emphasizing the need to examine variant read support on the ExAC browser. Furthermore, we know that truncating *ASXL1* variants commonly occur in individuals as they age (associated with CHIP), and it appears individuals with these variants skews older (median age 70). Also associated with pathogenic *ASXL1* variants is a mildly increased risk of developing MDS, myeloid malignancies, and even solid tumors (Artomov et al. 2016). The inclusion of exome data from TCGA may increase the prevalence of these variants, highlighting the need to use the non-TCGA ExAC data subset in some situations. Although based on the information currently available we were not able to conclusively attribute all truncating *ASXL1* variants in the ExAC database to technological artifacts or somatic mosaicism, we have presented a potential explanation for the presence of most (and perhaps all) such variants in reference population databases.

Although *ASXL1* provides an excellent illustration of how somatic mosaicism may confound the analysis of pathogenic variants, the issue is not unique to this gene. It is also worth considering *TET2* and *DNMT3A*, two other genes commonly mutated during clonal hematopoiesis (Genovese et al. 2014; Jaiswal et al. 2014). Germline variants in *TET2* are not associated with a known syndrome, however *de novo DNMT3A* variants cause autosomal dominant Tatton-Brown-Rahman syndrome, an overgrowth condition characterized by tall stature, macrocephaly, distinctive facial features, and intellectual disability (Tatton-Brown et al. 2014). *DNMT3A* encodes a DNA methyltransferase that is important in establishing DNA methylation patterns during development (Okano et al. 1999). Putative truncating variants in *DNMT3A* are present in ExAC and as with *ASXL1* demonstrate signs of somatic mosaicism (data not shown). However, the majority of cases of Tatton-Brown-Rahman syndrome have been attributed to pathogenic missense variants (Tatton-Brown et al. 2014).

One of the *DNMT3A* missense variants associated with Tatton-Brown-Rahman syndrome, p.Arg749Cys (Tatton-Brown et al. 2014), is reported in three individuals in ExAC. Based on allele balance (11%, 20%, and 24%), the presence of this pathogenic *DNMT3A* variant in ExAC can also likely be attributed to somatic mosaicism. The rounded ages of the two individuals with the 20% and 24% allele balance were 70, and the individual with the 11% allele balance was part of a cancer cohort for which age was not available. Another *DNMT3A* missense variant, p.Arg882His, has recently been associated with Tatton-Brown-Rahman Syndrome (Heeley et al. 2016; Kosaki et al. 2016), with the germline nature of this variant confirmed via buccal samples. The arginine residue at position 882 represents the most prevalent hotspot for pathogenic *DNMT3A* variants in acute myeloid leukemia (Ley et al. 2010; Lu et al. 2016), and the p.Arg882His variant was detected in 66 individuals (55 from non-cancer cohorts), a frequency of more than 1 in 1,000 individuals in ExAC. The p.Arg882His variant also demonstrates allele imbalance (the median allele balance is 19% with an IQR of 8-30% for the 55 individuals from non-cancer cohorts), which is consistent with somatic variants. Given the high burden of evidence required to consider a missense variant pathogenic (Richards et al. 2015), less well-characterized *DNMT3A* variants present in reference databases may be disregarded and diagnoses of Tatton-Brown-Rahman syndrome could be missed.

## DISCUSSION

Reference population databases such as ExAC are invaluable for characterizing the genetic diversity of human populations (MacArthur et al. 2014). These resources have enabled researchers and clinicians to better interpret variants as potentially pathogenic or benign based on allele frequency. However, these databases are known to contain individuals with pathogenic variants, including variants that cause recessive disorders, variants associated with incomplete penetrance, or variants associated with variable expressivity resulting in mildly affected individuals. Here, we present hematopoietic mosaicism as another caveat to consider in determining the pathogenicity of a variant based on its presence in a population database. When variants arise by somatic mutation, those that provide a growth advantage to hematopoietic cells can lead to clonal expansion resulting in these variants being present at a higher allele frequency than is typical for somatic variants. Such somatic variants with a relatively higher allele balance are more often mistakenly being called by variant calling algorithms that are designed to detect germline genetic variants.

The presence of any BOS-associated *ASXL1* variant in the ExAC database is unexpected for an autosomal dominant, severe, pediatric-onset condition that is assumed to be fully penetrant. Our analysis suggests that p.Arg404Ter is observed in individuals with allelic imbalance suggesting somatic mosaicism and clonal expansion. Most frequently these individuals were elderly or from a cancer cohort (both associated with CHIP). It is also possible that some individuals from non-cancer cohorts also have a past history of cancer or later develop cancer. In general, the *ASXL1* truncating variants (including another BOS-associated variant) in the ExAC database have variant allele imbalance consistent with hematopoietic somatic mosaicism.

Our results could also extend to genes associated with other autosomal dominant disorders. A recent study questioned the penetrance of pathogenic variants for intellectual disability and related disorders, including *ASXL1* (Ropers and Wienker 2015). The authors noted the difficulty of reconciling the apparent reduced penetrance of pathogenic *ASXL1* variants based on population database frequencies with the fact that inherited *ASXL1* variants have never been observed in BOS. Our study proposes that age-related somatic mosaicism with clonal expansion in the hematopoietic lineage could potentially account for these contradictory observations, and may even provide an explanation for other pathogenic variants identified in the Ropers and Weinker study. Awareness of the potential for somatic mosaicism is warranted when performing variant investigation for *ASXL1, DNMT3A,* and perhaps many other genes.

While our results suggest that somatic mosaicism with clonal expansion is a likely means by which some pathogenic variants are observed in non-syndromic individuals, we acknowledge that there are limitations to our study. We cannot discount the possibility that some of the variants excluded from our analysis (e.g. a subset of the p.Gly646TrpfsTer12 variants) are genuine pathogenic variants. Similarly, we have not demonstrated that all the truncating variants included in our analysis are pathogenic variants *in vitro* or *invivo*, though the identification of two previously published BOS-associated variants suggests that we are enriching for variants that impact ASXL1 function. We were not able to definitively confirm the somatic nature of the identified variants, as additional tissue samples were not available from the individuals in ExAC. Finally, we acknowledge that we could not explain all truncating *ASXL1* variants observed in ExAC, particularly those with normal allele balance in younger individuals from non-cancer cohorts. However, by attributing many of the truncating *ASXL1* variants to technological artifacts or somatic mosaicism, our analysis decreased the number of unexplained individuals with truncating *ASXL1* variants in ExAC from 395 to approximately 16. We believe that this study helps address the issue of pathogenic variants observed in population databases (Ropers and Wienker 2015), and will alleviate confusion during variant assessment using the current American College of Medical Genetics guidelines (Richards et al. 2015).

This investigation provides important guidance for variant analysis. In certain cases, one cannot dismiss the pathogenicity of a variant associated with an autosomal dominant condition based on the presence of that variant in reference population databases. Additional information regarding the individuals in the database may need to be examined, such as evidence for potential allelic imbalance, age, and whether they belong to a disease cohort such as patients diagnosed with cancer. Special attention should be paid to genes susceptible to the accumulation of age-related somatic variants in hematopoietic lineages, particularly as this is the source of DNA in most clinical genetic testing. Although we were fortunate to identify this issue due to our understanding of the role of *ASXL1* and *DNMT3A* in clonal hematopoiesis, this phenomenon might be overlooked in less well-characterized genes.

This work highlights the importance of certain features of the ExAC browser and suggests other areas where further annotation is needed. As it may relate to the interpretation of some variants, non-TCGA subset of the ExAC dataset is available for download. Additionally, a future goal is to include summarized age data on the browser. Examination of the raw sequence read data available on the ExAC browser is an important step of variant review, to identify any issues with the called variant, including skewed allele balance. Further work is needed to identify systematic deviations in allele balance in reference population databases, and it is a future goal to flag such variants in ExAC, although such an analysis would be not be straight-forward at present. In the mean time it is important that clinical laboratories and clinicians tasked with variant interpretation remain vigilant regarding the issue of somatic variants in all current reference databases.

The increasing volume of next-generation sequencing tests is already taxing a system that still largely relies on expert curation. Bioinformatics pipelines that filter out presumed benign variants are useful, but as our results have shown, over-dependence on variant frequencies in reference databases is dangerous, and establishing hard frequency thresholds for filtering will require careful consideration of complicating issues such as somatic mosaicism. As the trend toward automation continues it will be important not only to refine the rules of variant assessment, but also to recognize when the unique biology of certain genes may create important exceptions to those rules.

## Acknowledgements

We are grateful to the family for their willingness to share their daughter’s case with the medical community. We also thank the clinical genomics laboratory at ARUP for performing exome sequencing. We would like to thank the members of the Exome Aggregation Consortium for creating this resource. We appreciate the efforts of MacArthur lab members, particularly Monkol Lek, Konrad Karczewski, and Kaitlin Samocha for the development of resources for analysis of the ExAC dataset, several of which were very helpful in this work. We would also like to acknowledge John Carey, Oliver Tam, Wei Shen, Philippe Szankasi, Michael Van Ness, Tim Tidwell, and Alex Chapin for helpful discussions in Utah. C.M.C. is supported by The University of Utah Department of Pathology and ARUP Laboratories and A.H.O.-L. is supported by a Pfizer/ACMG Foundation Translational Genomic Fellowship. The ExAC analysis was supported by NIH grants NIGMS R01 GM104371 and NIDDK U54 DK105566.

## Author contributions

C.M.C., A.H.O.-L., T.T., D.G.M and R.M conceived and designed the study. H.R.U. evaluated the patient and contributed the clinical case description and patient photographs. T.T., C.M.C., and R.M. analyzed the patient’s exome sequencing data. A.H.O.-L., B.B.C., B.W., E.V.M., and D.P.B. collected the data from ExAC. A.H.O.-L. and C.M.C. analyzed and interpreted the variant frequency and allele imbalance. C.M.C. drafted the manuscript. C.M.C., H.R.U., T.T., A.H.O.-L., B.B.C., B.W., E.V.M., D.P.B., D.G.M. and R.M.critically revised the manuscript.

